# Synaptic Encoding of Time in Working Memory

**DOI:** 10.1101/2025.04.21.649874

**Authors:** Gianluigi Mongillo, Misha Tsodyks

## Abstract

The processing of temporally-extended sequences of stimuli critically relies on Working Memory (WM). Yet, how WM supports the encoding and retrieval of novel sequences is unknown. Existing theories rely on associative learning driven by repetitions and are, thus, unable to explain how people can reproduce novel sequences of stimuli immediately. Here, we propose that detailed temporal information about a novel sequence can be rapidly stored in WM by short-term synaptic plasticity over multiple time scales. To substantiate this proposal, we extend our previously-proposed synaptic theory of WM to include synaptic augmentation, besides more short-lived depression and facilitation, consistently with experimental observations. The long time scales associated with augmentation naturally lead to the emergence of a temporal gradient in the synaptic efficacies, which can be used to immediately replay, at normal speed or in a time-compressed way, novel sequences. The theory is consistent with behavioral and neurophysiological observations.

## 1 Introduction

Purposeful behavior requires storing and retrieving relevant information over multiple time scales. Typically, this information also includes a temporal component that is key to achieve the goal. For instance, to reach the closest coffee place we just asked directions to, we have to turn left at the next corner, walk one block, and then turn right. We’ll get no espresso following the directions in the *wrong* order.

The Working Memory (WM) – a specialized, low-capacity component of the memory system – is believed to be responsible for rapidly encoding and maintaining novel information (e.g., the directions we just asked) over short time scales [Cowan, 2001, Baddeley, 2003]. But, exactly, which information about a novel sequence of stimuli is stored in WM?

People effortlessly remember short but otherwise arbitrary (i.e., novel) sequences of familiar stimuli. For instance, people routinely hum short tunes they have just heard while musicians can even replay them with fidelity. A person can finger-tap, out of memory and with good accuracy, a pattern of irregularly spaced clicks spread over a few seconds which has been just experienced. The encoding of serial order, in particular, has been extensively investigated in the *serial recall* task [Kahana, 2012]. In this task, a list of randomly chosen items (e.g., words) is presented sequentially to the subject that, then, has to recall them in the presented order. This task is thought to rely on WM and, indeed, the number of correctly recalled items – typically about 4 items – is a standard measure of WM capacity. Interestingly, people almost invariably recall short lists (i.e., within WM capacity) in the presented order, even without explicit instructions to do so, as in the *free recall* task [Dimperio et al., 2005, Ward et al., 2010, Grenfell-Essam and Ward, 2012].

These observations suggest that WM rapidly and automatically stores quite detailed temporal information about a novel sequence of familiar stimuli, in addition to information about the stimuli themselves.

The models originally proposed for the computational architecture of WM have no mechanism for the encoding of temporal information [Cowan, 2001, Baddeley, 2003]. As to neuronal models of WM (i.e., short-term memory maintenance), there have been different proposals. The most popular idea is that active maintenance relies on the co-existence of stable steady states of activity in the memory network (attractors) that are selected by stimulus presentations [Amit, 1995, Amit and Brunel, 1997, Wang, 2021]. The current state of activity, thus, reflects the recent history of stimulation. This mechanism can store the identity of the stimuli in the sequence but not information about their relative timing (e.g., the order of occurrence); this information would then need to be learned in the course of multiple repetitions of the *same* sequence [Kleinfeld, 1986, Sompolinsky and Kanter, 1986].

Partly to address the inability of attractor networks to *rapidly* store temporal information, an alternative account has been proposed [Maass et al., 2002, Buonomano and Maass, 2009]. In this account, active maintenance relies on the transitory, but high-dimensional, responses elicited by stimulus presentation in the memory network (liquid state machine). Such a mechanism can rapidly store the identity of the stimuli as well as detailed information about their times of occurrence, thanks to the high-dimensionality of the response. However, it is now the read-out of this information that needs to be learned, again in the course of multiple repetitions [Cueva et al., 2020, Zhou et al., 2023].

Somehow surprisingly in view of their profound differences, these two accounts make one identical prediction; temporal information about a *novel* sequence of stimuli is not immediately available (e.g., to produce some behavior), either because it has not been stored yet (attractor networks) or because it cannot yet be read-out (liquid state machines). We have just discussed evidence contrary to this prediction.

We have proposed that information is maintained in WM by synaptic facilitation within the neuronal populations that code for the items, rather than by the enhanced, persistent activity of those populations [Mongillo et al., 2008]. Facilitation is an experimentally well-characterized transient enhancement of the synaptic efficacy that is quickly induced by pre-synaptic spiking activity and can last for up to several seconds [Markram et al., 1998, Zucker and Regehr, 2002]. In particular, facilitation was reported at inter-pyramidal connections in the prefrontal cortex, a region heavily implicated in WM [Hempel et al., 2000, Wang et al., 2006]. The theory is compatible with multiple experimental observations and motivated further experiments aimed at disentangling persistent activity and information maintenance [Rose et al., 2016, Wolff et al., 2017, Panichello et al., 2024].

In the framework of the synaptic theory of WM, the maintenance of information can be achieved by different regimes of neuronal activity, depending on the background input to the network; at increasing levels of the background input, these regimes are: (i) activity-silent, where the information is transiently maintained without enhanced spiking activity; (ii) low-activity, where the information is periodically refreshed, at low rate, by brief spontaneous reactivations of corresponding neuronal populations (i.e., population spikes, PSs); (iii) persistent-activity, where the information is maintained by tonically active neuronal populations.

Facilitation is not the only form of transient synaptic enhancement induced by repetitive pre-synaptic activity. Experiments reveal other forms, such as augmentation and potentiation, which build up more slowly than facilitation but are significantly more long-lived [Fisher et al., 1997, Thomson, 2000, Fioravante and Regehr, 2011]. As a result, the instantaneous value of the synaptic efficacy can reflect the history of pre-synaptic activation over tens of seconds (i.e., the time scale of augmentation) or even minutes (i.e., the time scale of potentiation) rather than just seconds (i.e., the time scale of facilitation). In the present contribution, we propose that such a transient synaptic enhancement over multiple time scales allows the encoding of *both* stimulus *and* temporal information in the instantaneous synaptic efficacies.

To substantiate this proposal, we extend the synaptic theory of WM to include synaptic augmentation, observed in the prefrontal cortex at the same synapses that exhibit significant short-term facilitation [Hempel et al., 2000, Wang et al., 2006].

## 2 Results

To illustrate the putative role of synaptic augmentation in the encoding of temporal information, we consider the simplified setting used in [Mi et al., 2017]. The network is composed of *P* distinct excitatory populations, that represent the memory items, and one inhibitory population, that prevents simultaneous activity at enhanced rates in the excitatory populations. The recurrent synaptic connections within each excitatory population display short-term synaptic plasticity according to the Tsodyks-Markram (TM) model [Markram et al., 1998]. The population-averaged synaptic input to population *a* (*a* = 1, …, *P*), *h*_*a*_, evolves in time according to

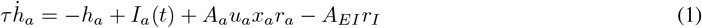

where *τ* is the neuronal time constant; *I*_*a*_(*t*), the external input to population *a*, is the sum of two components: a background input, to control the activity regime of the network, and a selective input, to elicit enhanced activity during the presentation of the corresponding item; *A*_*a*_ is the average strength of the synapses within excitatory population *a*; *r*_*a*_, the average activity of population *a*, is a smoothed threshold-linear function of *h*_*a*_, i.e.,

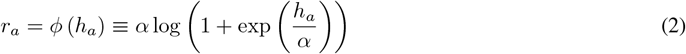

where *α >* 0 is a parameter controlling the smoothing; *u*_*a*_ and *x*_*a*_ are, respectively, the levels of short-term facilitation and depression of the recurrent synapses within population *a*; *A*_*EI*_ is the strength of the synapses from the inhibitory population to any excitatory population; *r*_*I*_ = *ϕ* (*h*_*I*_) is the average activity of the inhibitory population, and

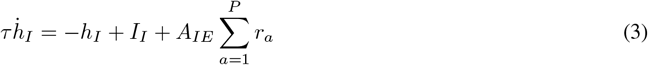

where *I*_*I*_ is the constant background input to the inhibitory population and *A*_*IE*_ is the strength of the synapses from any excitatory population to the inhibitory population.

The levels of short-term facilitation and depression, *u*_*a*_ and *x*_*a*_, evolve in time according to [Tsodyks et al., 1998]:

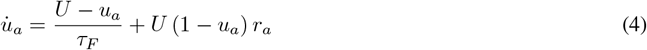

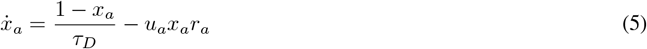

where *U* is the baseline release probability; *τ*_*F*_ and *τ*_*D*_ are the facilitation and depression time constants, respectively. In words: Activity in the population induces both facilitation, i.e., it increases *u*_*a*_, and depression, i.e., it decreases *x*_*a*_, while, in the absence of activity (i.e., *r*_*a*_ = 0), facilitation and depression decay to their respective baseline levels, *u*_*a*_ = *U* and *x*_*a*_ = 1.

In [Mi et al., 2017], the *A*_*a*_’s in Equation (1) are time-independent parameters with the same value for all the excitatory populations. By contrast here, to model synaptic augmentation, the *A*_*a*_’s are activity-dependent dynamic variables that increase with the *r*_*a*_’s according to

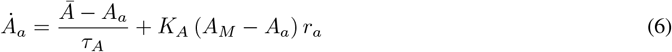

where 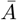 is the basal strength of the synapses within an excitatory population observed when there is no presynaptic activity (i.e., *r*_*a*_ = 0); *τ*_*A*_ is the augmentation time constant, *K*_*A*_ controls how fast the average strength of the synapses, *A*_*a*_, increases with the activity, and *A*_*M*_ is the maximal synaptic strength that can be induced by augmentation. In the following, we take 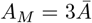.

The physiological mechanisms responsible for synaptic augmentation are poorly understood [Fisher et al., 1997, Thomson, 2000, Fioravante and Regehr, 2011]. Equation (6) provides a minimal phenomenological description of synaptic augmentation in the spirit of the original TM model [Markram et al., 1998]. However, as it will become clear in the following, our results do not depend critically on this modeling choice. For instance, one would obtain the same results by modeling augmentation as an activity-dependent increase in the baseline release probability *U* (data not shown). Facilitating synaptic transmission observed at inter-pyramidal synapses in the prefrontal cortex is well described by the above model with the following choice of synaptic parameters: *U* ~ 0.2, *τ*_*F*_ ~ 1s, *τ*_*D*_ ~ 0.1s, *τ*_*A*_ ~ 10s and *K*_*A*_ ≪ 1 [Hempel et al., 2000, Wang et al., 2006, Barri et al., 2016]. The full set of network and short-term plasticity parameters used in the simulations can be found in the caption of Fig. 1.

**Figure 1.**
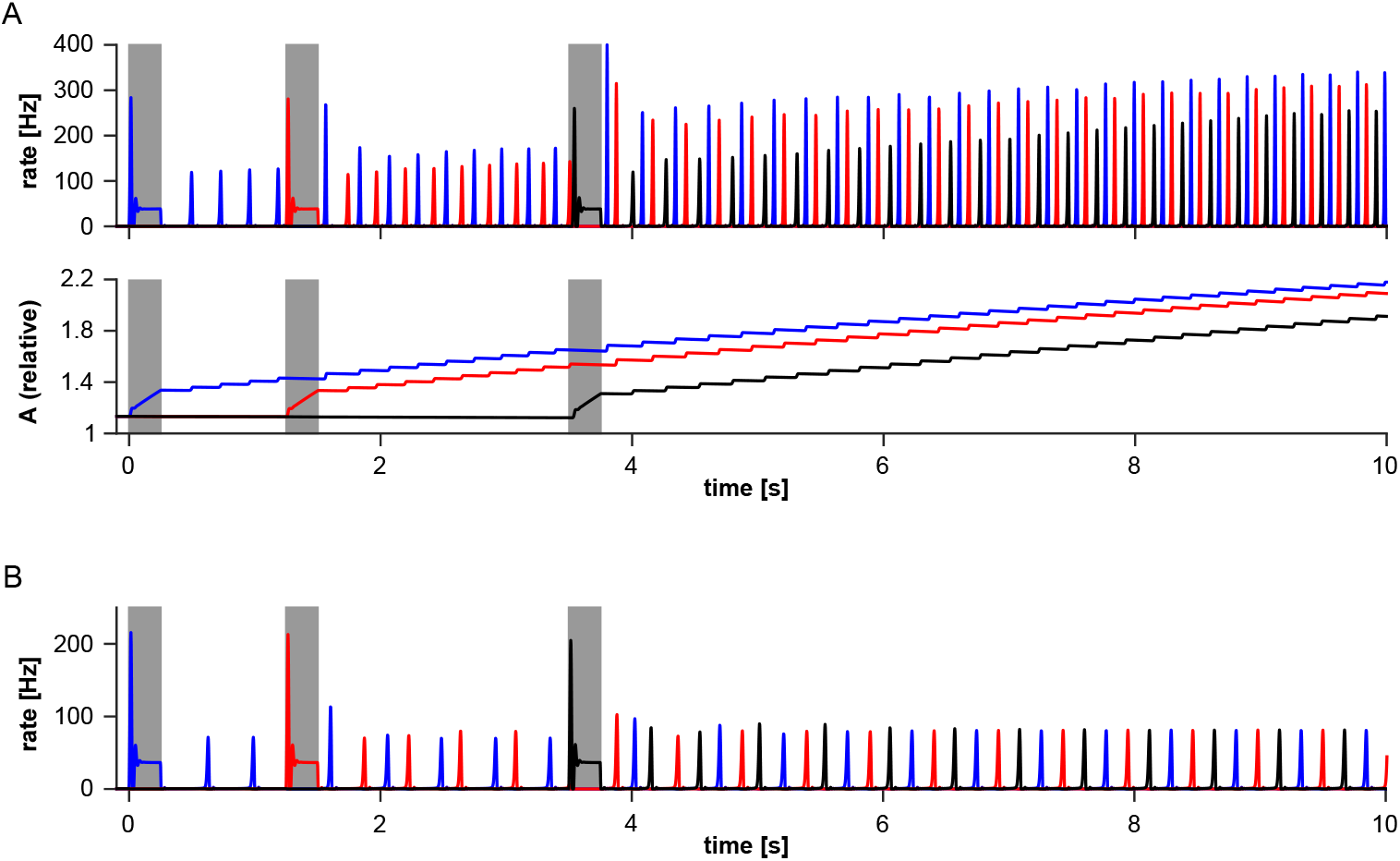
Network activity encodes sequence information. Network responses to 3 sequentially presented items with (A) and without synaptic augmentation (B). The bottom panel in (A) shows the level of synaptic augmentation in the corresponding synaptic populations. The presentation of an item is simulated by a 10-fold increase of the background input selectively to the corresponding neuronal population for 250ms (gray areas). The background input to the remaining populations is kept constant at its baseline level. Network parameters: *P* = 16, *τ* = 8ms, *α* = 1.5Hz, *A*_*EE*_ = 6.0, *A*_*EI*_ = 1.1, *A*_*IE*_ = 1.75, *I*_*bkg*_ = 8.0Hz; Short-term plasticity parameters: *U* = 0.3, *K*_*A*_ = 0.01, *τ*_*D*_ = 0.3s, *τ*_*F*_ = 1.5s, *τ*_*A*_ = 20s.

In Fig. 1A, we show the response of the model network to a sequence of 3 items with variable inter-item intervals. The interval between the onset of the first and second item is 1 second, while the interval between the onset of the second and the third item is 2 seconds. Following the presentation of the last item, the neuronal populations that have been stimulated reactivate in a repeating cycle, indicating that the corresponding items are being actively maintained in WM. Importantly, this regime of activity does not correspond to a steady state of the network dynamics. This is evident from the amplitudes of the PS and from the levels of synaptic augmentation in the reactivating neuronal populations (Fig. 1A, bottom panel) that are still changing with time. Note that the amplitudes of the PS are different for the different populations.

For comparison, we show in Fig. 1B the response of the network to the same sequence in the absence of synaptic augmentation (i.e., 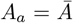 and *K*_*A*_ = 0). In this case the network dynamics rapidly converge to a steady, attractor state (on a time scale ~ *τ*_*F*_); the amplitude of the PS is the same for all the reactivating populations. Once this state is reached, the network carries no information about the sequence beyond the identity of the stimuli composing the sequence.

The transient regime exhibited by the model network in the presence of augmentation is long-lived because the level of augmentation grows slowly with neural activity. As can be seen in the bottom panel of Fig. 1A, significant augmentation only occurs during the reactivations. The increase of the augmentation level with each reactivation is mainly controlled by *K*_*A*_ and *K*_*A*_ is small, consistently with the experiments. As a result, the levels of synaptic augmentation in the reactivating neuronal populations are still changing with time long after the presentation of the last item in the sequence. At the same time, the decay of the level of augmentation between two consecutive reactivations of the same population ( ~ *τ*_*D*_) is negligible, because *τ*_*D*_ ≪ *τ*_*A*_. Therefore, the longer an item has been active in WM – that is, the larger the number of reactivations – the larger the corresponding level of augmentation. Indeed, the level of augmentation encodes, quite accurately, the time elapsed since item’s presentation (Fig. 1A, bottom panel).

If augmentation was the only form of synaptic plasticity present in the network, the encoding of an item in WM would require long presentation times, or alternatively high firing rates upon presentation, precisely because *K*_*A*_ is small. Instead, rapid encoding is made possible by the presence of the short-term facilitation, which builds up significantly faster than augmentation, as *U* ≫ *K*_*A*_. For the same reason, however, the level of facilitation rapidly reaches the steady state; therefore, short-term facilitation alone is unable to encode temporal order (see Fig. 1B). Thus, our model requires the existence of transitory synaptic enhancement on at least two time scales, such that longer decays are accompanied by slower build-ups. Intriguingly, this pattern is experimentally observed [Fisher et al., 1997].

In summary, in the presence of synaptic augmentation, WM activity naturally encodes the temporal structure of a novel sequence of familiar items, besides encoding information about the identity of the single items. As this information is present in the levels of synaptic augmentation and in the amplitudes of the PSs during the reactivations, it is readily accessible to a downstream read-out network.

To illustrate this important point, we consider a simple read-out mechanism to reconstruct/replay the sequence stored in WM (Fig. 2A). Each item-selective population in the memory network provides excitatory inputs to the corresponding item-selective population in a read-out network. For simplicity, we assume that the excitatory synapses between the memory and the read-out network exhibit the same dynamics as the excitatory synapses within the memory network. The activation of a population in the read-out network signals the retrieval of the corresponding item. The item-selective populations in the read-out network do not interact with each other. Rather, they receive a uniform background input (i.e., the same for all populations) that effectively sets the threshold for their activation. To read-out the contents of WM, following the presentation of the last stimulus, the background input ramps up until the first population in the read-out network activates and then remains constant.

**Figure 2.**
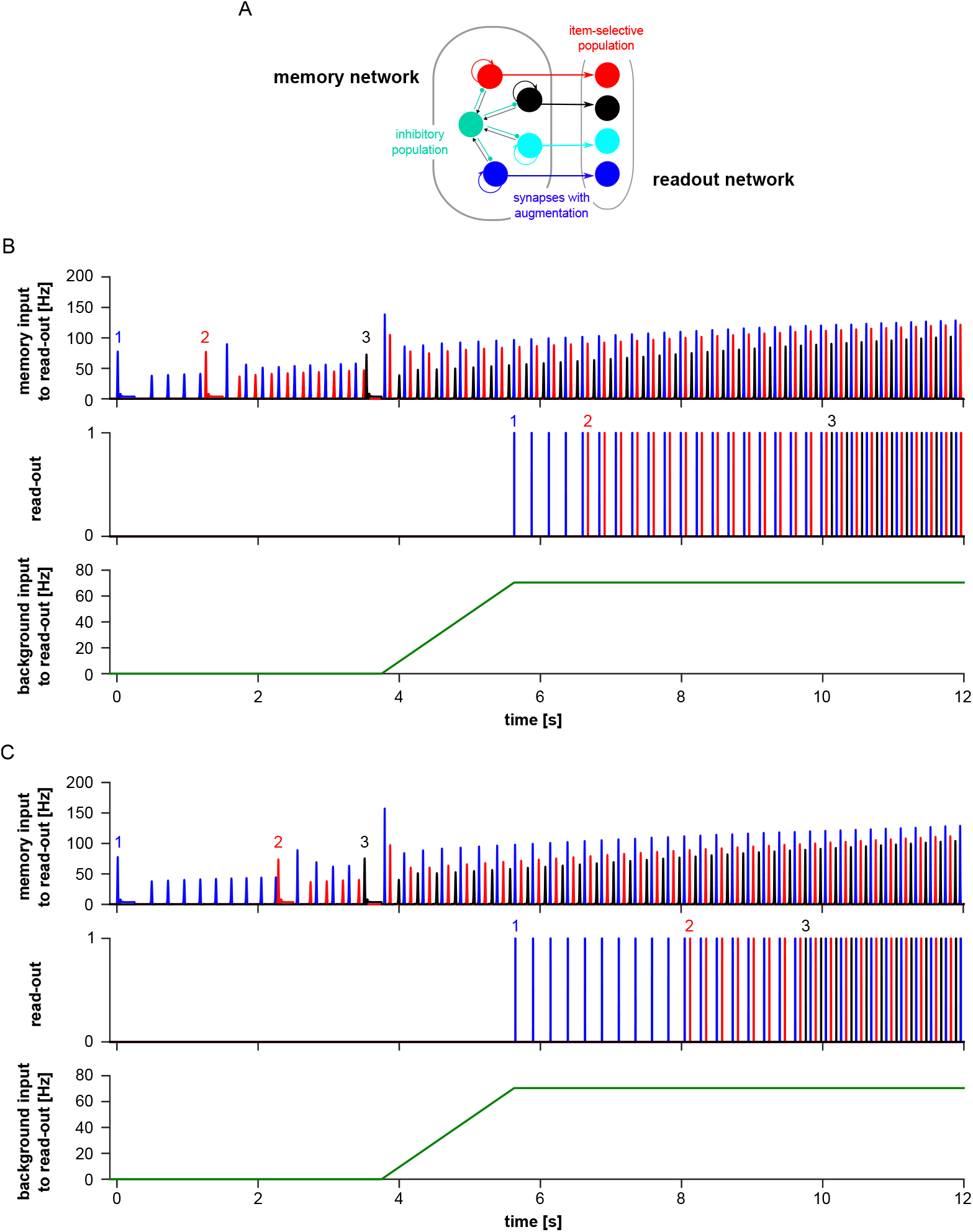
Replay of a novel sequence. (A) Architecture of the memory and read-out network (see main text for details). (B) Top panel: Input to the read-out network from the memory network. Middle panel: Activation state of the item-selective populations in the read-out network, as determined by comparing the sum of the input from the memory network and of the background input to the threshold. Bottom panel: Time course of the background input to the read-out network. (C) Same as (B) for a different sequence. Note that the time course of the background input to the read-out network is the same in (B) and (C).

This mechanism results in an approximate replay of the sequence stored in WM. We illustrate this in Fig. 2B and C for two different sequences and the *same* read-out network. This is because the temporal evolution of the level of augmentation and of the amplitude of the PSs in the active populations are approximately time-translation invariant; that is, they are approximately the same when aligned to stimulus onset. Hence, the inputs from the memory to the read-out network will reach the same level (i.e., the same threshold) at time intervals that (approximately) match the time intervals between the presentations of the corresponding items. For the same reason, the accuracy of the replay is rather robust against (reasonable) changes in the rate of increase of the input to the read-out network.

The augmentation gradient can also be used to fast-replay the sequence stored in WM. Fast replay has been suggested as a mechanism for consolidating the storage of information in the long-term memory, by bringing patterns of neuronal activity representing temporally distant events within a time window in which long-term synaptic plasticity can most effectively operate [Melamed et al., 2004, Jensen and Lisman, 2005]. The time-compressed replay of the sequence is initiated by decreasing the level of background input to the WM network for a time ~ *τ*_*F*_ (Fig. 3, bottom panel, downward arrow). This prevents further reactivations (Fig. 3, top panel) and the corresponding synaptic variables start decaying toward their baseline levels (Fig. 3, middle panel). The background input is then raised again to a suitably larger level (Fig. 3, bottom panel, upward arrow). In the time interval where reactivations are suppressed, the levels of augmentation, i.e., the *A*_*a*_’s, do not change significantly because *τ*_*F*_ ≪ *τ*_*A*_. However by the end of the same time interval, short-term depression and facilitation variables will be close to their corresponding baseline levels (i.e., *x*_*a*_ ≃ 1 and *u*_*a*_ ≃ *U* for *a* = 1, …, *P*). When the background input is raised above a critical level, the steady, low-rate state of activity becomes unstable for the once-active neuronal populations. Therefore, they will start reactivating, with the most unstable one (i.e., the one with the larger *A*_*a*_) reactivating first, the next most unstable one reactivating second, and so on [Mi et al., 2017], hence replaying the sequence in the order of stimuli presentation (Fig. 2, top panel). Note that the network replays the sequence in about 250 ms, thus achieving a compression factor of about 10.

**Figure 3.**
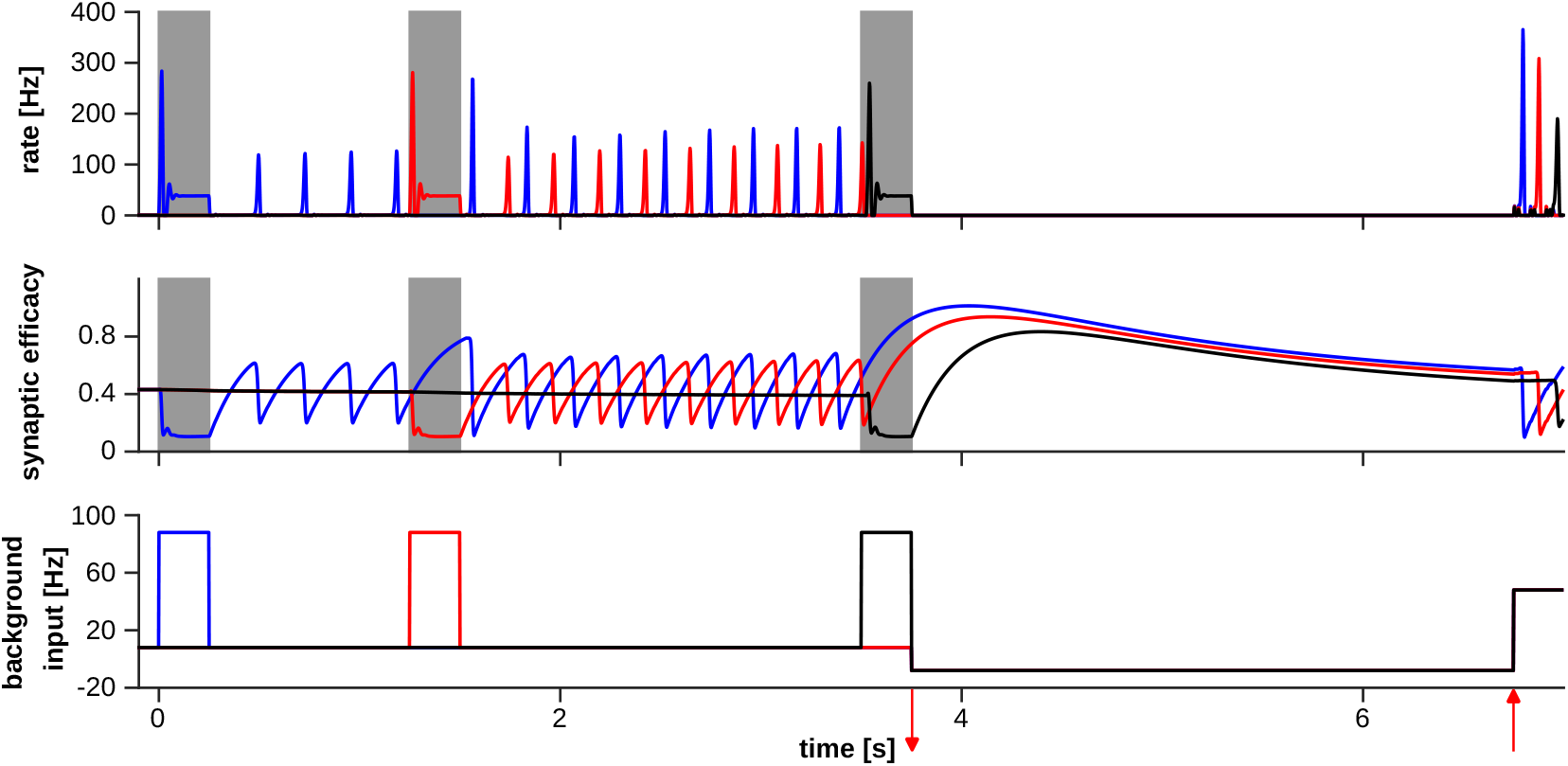
Fast-replay of a novel sequence. The top panel shows the response of the network to the external inputs depicted in the bottom panel. After the presentation of the sequence, the background input is first decreased (downward red arrow in the bottom panel) for 3 seconds and then increased (upward red arrow in the bottom panel) for 250 milliseconds. The middle panel shows the resulting time course of *Aux* in the corresponding synaptic populations. Immediately before the background input is increased again, *Aux* ≃ *AU*.

There is significant experimental evidence that items can be maintained in WM in different *representational* states and that these states can be rapidly altered by task demand [LaRocque et al., 2014, Oberauer and Awh, 2022]. A case in point is the study of [Rose et al., 2016], who used a retro-cue design to manipulate these putative representational states (see Fig. 4A). Briefly, human subjects performed a two-item delayed recognition task, with two retro-cues and two recognition probes per trial. Following the presentation of the items and a delay period, the first retro-cue informed the subject about which of the two items will be probed in the impending recognition test, after a delay period. The cued item is considered *prioritized* for upcoming behavior. After the first recognition test, a second retro-cue indicated the item to be probed in the second recognition test, following another delay period. Importantly, each retro-cue randomly prioritized either of the two items with the same probability and, hence, both items had to be kept in WM until the second retro-cue. Rose et al. [2016] found that, during the initial delay period, both items could be reliably decoded from the fMRI signal. By contrast, during the delay period between a retro-cue and the subsequent recognition test, only the prioritized item could be reliably decoded. However, the decodability of the de-prioritized item was recovered by transcranial magnetic stimulation.

**Figure 4.**
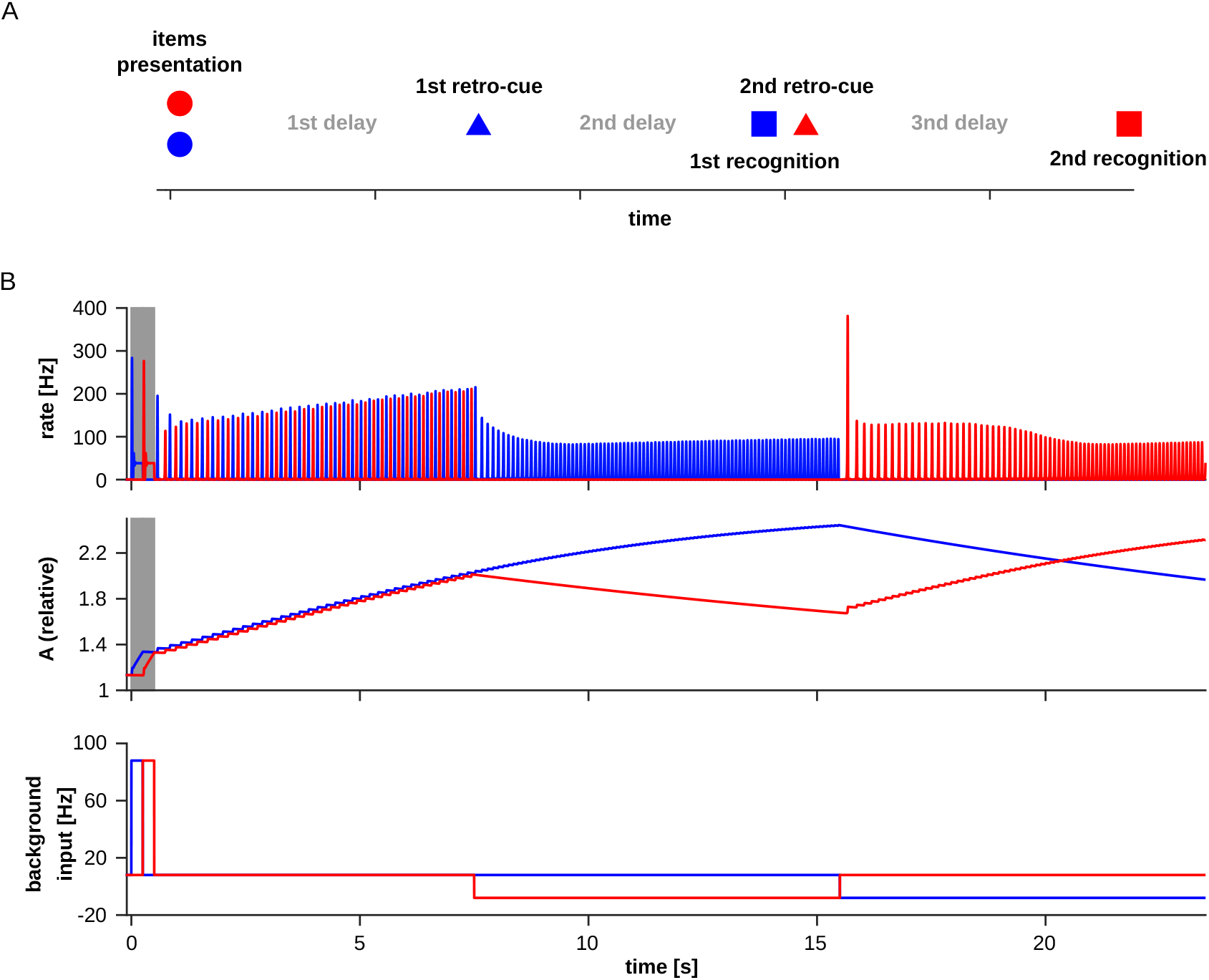
Reactivation of an activity-silent memory. (A) Cartoon illustrating the task events in the study of [Rose et al., 2016]. Circles indicate items presentation, triangles indicate cue instructions, squares indicate recognition tests. In this example, the first retro-cue prioritizes the ‘blue’ item, that is then probed during the first recognition test, and the second retro-cue prioritizes the ‘red’ item, which is probed in the second recognition test. See main text and [Rose et al., 2016] for details. (B) The top panel shows the response of the network to the external inputs depicted in the bottom panel. 8 seconds after the presentation of the two items, the background input to the neuronal population selective to second item (red) is decreased for 8 seconds, and then restored to its original level, while decreasing the background input to the neuronal population selective to the first item (blue). The middle panel shows the resulting time course of the level of augmentation in the corresponding synaptic populations.

The experimental observations of [Rose et al., 2016] (see also [Wolff et al., 2017]) suggest that there are at least two different states of ‘maintenance’ in WM with distinct neurophysiological signatures. Though the relationship between the fMRI signal and neural activity is not an obvious one, these results have been interpreted as indicating that these two states differ in their level of neural activity. In particular, the failure to decode would indicate the absence of enhanced spiking activity (but see [Barbosa et al., 2021]).

Our model network can reproduce these experimental observations, as illustrated in Fig. 4B. Similarly to the experiment of [Rose et al., 2016], two items are presented and, after the initial delay period, one of the two is prioritized. We simulate the effect of the retro-cue on neural activity by assuming that the de-prioritized item receives a lower external input as compared to the prioritized item (Fig. 4B, bottom panel). Following the first retro-cue, the neuronal population encoding the prioritized item keeps reactivating while the neuronal population encoding the de-prioritized item stops reactivating (Fig. 4B, top panel). We account for the results of [Rose et al., 2016] by assuming that only reactivating items can be decoded.

In the absence of neural activity, the ‘working memory’ of the de-prioritized item is kept by the augmentation level (Fig. 4B, middle panel). To illustrate this, we prioritize the previously de-prioritized item with the second retro-cue and, indeed, the corresponding neuronal population resumes its reactivating dynamics (Fig. 4B, top panel). In the presence of synaptic augmentation, the maximal time span of the activity-silent regime is controlled by *τ*_*A*_, which is of the order of 10 seconds, which is consistent with the experimental observations of [Rose et al., 2016]. We note that in the model originally proposed in [Mongillo et al., 2008], the time span of the activity-silent regime is limited by *τ*_*F*_ ( ~ 1 s), which is not consistent with the results of [Rose et al., 2016].

## 3 Discussion

We propose that transient, non-associative synaptic plasticity over multiple time scales can support the temporally-structured encoding of a sequence of stimuli. We have illustrated this idea in a minimal model network that extends the synaptic theory of WM to include synaptic augmentation, besides synaptic depression and facilitation. Our model allows the storage and the retrieval of short sequences of items by relying on synaptic plasticity mechanisms that are well-characterized experimentally, that is, the transient enhancement on multiple time scales of the synaptic efficacy driven *solely* by pre-synaptic activity [Fisher et al., 1997, Thomson, 2000, Fioravante and Regehr, 2011]. In the low-activity regime, where items are maintained by short-lived reactivations of the corresponding neuronal populations, the presence of synaptic augmentation naturally leads to a temporal gradient in the synaptic efficacies that encodes both the items and their relative times of occurrence. This gradient can then be used to replay the sequence either at normal speed or in a time-compressed way. The mechanism that generates the temporal gradient is robust, because it relies on the order-of-magnitude differences between the build-up and the decay time of the augmentation and those of the depression and facilitation.

A key prediction of our theory is that items are maintained in a low-activity regime. Indeed, if the items are maintained either in an activity-silent regime or by persistent, high firing rates, the proposed mechanism fails. In the first case, because in the absence of reactivations the gradient does not build up; in the second case, because the augmentation levels quickly saturate due to the high firing rates. This prediction is consistent with multiple experimental observations [Siegel et al., 2009, Fuentemilla et al., 2010, Lundqvist et al., 2016, Panichello et al., 2024, Liebe et al., 2025].

In multi-item working memory tasks, neural activity during the maintenance period is characterized by short episodes of spiking synchrony, detected as brief gamma bursts in the local field potential [Siegel et al., 2009, Lundqvist et al., 2016] or in the MEG/EEG signal [Fuentemilla et al., 2010]. Importantly, during a given gamma burst, only one of the items can be reliably decoded [Fuentemilla et al., 2010, Lundqvist et al., 2016], suggesting that the items are reactivated briefly and sequentially (i.e., one at a time) during maintenance. More direct support to this interpretation comes from recent electrophysiological studies [Panichello et al., 2024, Liebe et al., 2025]. By recording large neuronal populations ( ~ 300) simultaneously in the prefrontal cortex of monkeys performing a WM task, [Panichello et al., 2024] found that, during the maintenance period, the decoding of the actively held item from neural activity was ‘intermittent’; that is, decoding was only possible during short epochs ( ~ 100ms) interleaved with epochs (also ~ 100ms) where decoding was at chance level. The inability to decode resulted from a loss of selectivity at the population level, with a return of the single-neuron firing rates to their spontaneous (pre-stimulus) activity levels. The transitions between these two activity states (decodable/not-decodable) were coordinated across large populations of neurons in PFC. By recording single-neuron activity in the medial temporal lobe of humans performing a sequential multi-item WM task, [Liebe et al., 2025] found that during maintenance, neurons coding for a given item tended to fire at a specific phase of the underlying theta rhythm, again suggesting that the corresponding neuronal populations reactivate briefly and sequentially. In summary, these experimental results suggest that active memory maintenance relies on brief reactivations of the neural representations of the items, which we identify with the population spikes in our model, and that these reactivatations occur sequentially in time, as predicted by our theory.

We note that the proposed mechanism would still work if the items were maintained by tonically-enhanced firing rates, instead of population spikes, provided that those firing rates were suitably low. However, obtaining low firing rates in model networks of persistent activity is quite difficult.

Behavioral data in serial recall tasks strongly support the idea that serial order encoding is based on a *primacy gradient* [Grossberg, 1978, Farrell and Lewandowsky, 2004, Hurlstone and Hitch, 2015, 2018]. Our theory makes an explicit proposal as to its neurophysiological substrate: The primacy gradient is encoded by augmentation levels emerging due to interplay between synaptic and neuronal dynamics (as described above). As such, the generation of the gradient is an inescapable consequence of the active maintenance of an item in WM. This would naturally explain why the recall order tends to be the same as the presentation order also in free-recall tasks, provided that the sequence does not exceed WM capacity.

In the behavioral context, our theory also makes novel predictions. For example, the temporal gradient builds up gradually with the reactivations of the corresponding neuronal populations between consecutive presentations. This requires a presentation rate that is slow enough for these reactivations to occur in sufficient numbers. Hence, as the presentation rate is increased, the theory predicts that encoding of the serial order should degrade. Consistently with this prediction, increasing the presentation rate results in a larger number of transposition errors, that is, some items are recalled at the wrong serial position (see, e.g., [Farrell and Lewandowsky, 2004]). Experiments with very rapid serial visual presentation (RSVP) show that subjects cannot report the correct presentation order, even when the number of items is below capacity [Reeves and Sperling, 1986]. At the other extreme, if the presentation rate is too slow, or the list is too long, then the primacy gradient will also degrade either because of the saturation of synaptic augmentation or failure to actively maintain all the items. Consistent with this prediction, the spontaneous tendency to recall the items with the presented order in free recall tasks rapidly degrades with increasing list lengths [Grenfell-Essam and Ward, 2012].

To illustrate our theory in a simple setting, we used a minimal model network that neglects many physiological details. This, however, constitutes a limitation of the present study. It would be reassuring to see that the mechanism we propose here is robust enough to reliably operate also in spiking networks, in the presence of heterogeneity in both single-cell and synaptic properties. While we are fairly confident that this is the case, a spiking implementation of our model is beyond the scope of the present study and will be addressed in the future. Also, because of the simplicity of the model network, a comparison between the model behavior and the electrophysiological observations cannot be completely direct. Nevertheless the model qualitatively accounts for a diverse set of experimental data.

The model naturally generates ramping activity as a consequence of active maintenance, that is, the average level of activity in a reactivating neuronal population increases with time (see, e.g., Fig. 1). Ramping activity has been indeed proposed as a potential neuronal mechanism to encode time and it is commonly observed in electrophysiological studies of WM. A case in point is the recent study of [Cueva et al., 2020], who found ramping activity during the maintenance period in different delayed response tasks, regardless of whether timing was relevant for the task.

In neurophysiological studies of WM for sequences the conjunctive coding of item identity and serial-order information at the single-neuron level has been observed [Barone and Joseph, 1989, Funahashi et al., 1997, Xie et al., 2022]. Conjunctive coding refers to the modulation of neuron’s activity by both item and order information, so that, for instance, the average firing rate of the neuron during the delay period following different sequences with the same item changes depending on the position of the item in the sequence [Xie et al., 2022]. In our model the firing rate of a neuron is naturally sensitive to the temporal order due to the augmentation gradient (Fig. 2B). It remains to be seen whether our model in a more physiologically detailed setting can quantitatively account for some features of conjunctive coding as observed in experiments [Xie et al., 2022]. In this respect, an important caveat is that animals in these studies have been extensively trained on the task with a limited number of sequences. Extensive training and sequences’ repetition could lead to the emergence of stimulus-adapted neuronal representations via associative plasticity mechanisms [Botvinick and Watanabe, 2007, Gillett et al., 2020, Ryom et al., 2021].

The long time scales brought about by synaptic augmentation significantly extend the time span of memories maintained in the activity-silent state. As discussed in the Results section, the time scales of synaptic augmentation are fully compatible with the experimental observations, such as those of [Rose et al., 2016], suggesting that a memory that has been *silent* for ~ 10 seconds can still be retrieved upon cuing (see Fig. 4). Accordingly one would expect a large *storage* capacity in the activity-silent mode. For instance, by assuming that an item is to be refreshed every 10 seconds to prevent its loss (based on [Rose et al., 2016]), and that refreshing takes 100 milliseconds (based on [Panichello et al., 2024]), one would estimate a storage capacity of 100 items for the activity-silent WM. This raises the possibility that classical WM capacity, experimentally estimated with uncued *recall*, could result from the inability to retrieve the information, rather than from the inability to encode and/or maintain it. In this scenario, WM capacity is ultimately determined by the degree of *selectivity* that the background control – that we identify with the “central executive” or the “focus of attention” of cognitive theories – can attain.

## 4 Acknowledgements

G.M. work is supported by grants ANR-19-CE16-0024-01 and ANR-20-CE16-0011-02 from the French National Research Agency and by a grant from the Simons Foundation (891851, G.M.). M.T. is supported by the Israeli Science Foundation grant 1657/19 and Foundation Adelis.

